# Melanocyte-secreted Fibromodulin constrains skin inflammation in mice injected with lupus serum

**DOI:** 10.1101/2022.05.13.491848

**Authors:** Marianna Halasi, Abraham Nyska, Limor Rubin, Yuval Tal, George C. Tsokos, Irit Adini

**Affiliations:** Harvard Medical School, Department of Surgery, Center for Engineering in Medicine & Surgery, Massachusetts General Hospital, 51 Blossom Street, Boston, MA, 02114.; Sackler School of Medicine, Tel Aviv University, and Consultant in Toxicologic Pathology, Ramat Aviv 69978, Israel.; Allergy and Clinical Immunology Unit, Department of Medicine, Hadassah Medical Organization, Faculty of Medicine, Hebrew University of Jerusalem, Israel.; Harvard Medical School, Department of Medicine, Rheumatology, Beth Israel Deaconess Medical Center, E/CLS, 330 Brookline Ave, Boston, MA 02115.

## Abstract

Skin pigmentation has been linked to the development, prevalence, and severity of several immune-mediated diseases such as SLE. Here, we asked whether fibromodulin (FMOD), which is highly expressed in skin with light complexion, can explain the known differences in the magnitude of inflammation. C57 mice with different levels of pigmentation and FMOD were injected with human lupus serum to induce skin inflammation. Histopathologic studies revealed that black C57 FMOD+/+ that produce low levels of FMOD and white C57 FMOD -/- mice develop more severe inflammation compared with white FMOD +/+ mice. This study also revealed that dark pigmentation and FMOD deletion correlates with the increased numbers of Langerhans cells. Altogether, we identify low pigmentation and FMOD are linked to low severity of inflammation and approaches to promote FMOD expression should offer clinical benefit.

## INTRODUCTION

Epidemiologic studies underscore the importance of race in the population-specific inflammation incidence, which depend upon skin pigmentation levels. Systemic lupus erythematosus (SLE) is a chronic autoimmune disease, wherein ethnicity is a key factor in determining severity.^1^ People with darkly pigmented skin experience 2.4-fold higher incidence and prevalence rates compared to Caucasians.^2–4^ Although SLE is rare in Africa, it is more frequent among African descendants around the globe, suggesting the contribution of genetic and environmental factors. SLE patients of African descent, have a higher risk of acute onset, discoid rash, moreover, individuals with dark skin tones have a lower probability of achieving remission.^5–7^

Black/brown individuals differ from Caucasians in melanin production as well as melanocyte structure and function, despite the fact that both groups having a comparable number of melanocytes.^8^ UV radiation from the sun not only stimulates melanocytes to produce melanin, but it also activates skin-resident T helper lymphocyte cells through the epidermal antigen-presenting dendritic cells (DCs). Thus, epidermal-Langerhans cells (LCs; a subset of immature DCs that reside in the epidermis), and the inflammatory DCs (epidermal and dermal DCs) are thought to play a central role in the pathogenesis of skin flares.^9–12^ The skin resident DCs constantly survey the environment for pathogenic activity, including microbial invasion and injury. Immature DCs (imDC), when activated migrate through the basement membrane, during which they interact directly with extracellular matrix (ECM) microenvironment proteins, such as fibromodulin (FMOD), leading to their maturation and subsequent stimulation of the immune system.

We have previously shown that lowly pigmented (white) melanocytes secrete high levels of FMOD, which in return increases TGF-β1 secretion, a known inhibitor of DC maturation.^13–16^ Here, we present evidence that darker pigmentation contributes to the severity of skin inflammation in response to human SLE serum injection. Furthermore, we demonstrate that melanocyte secreted FMOD plays an important role in the serum induced skin inflammation.

## METHODS

### Mice

C57BL/6J (FMOD+/+) and mutant B6(Cg)-*Tyr^c-2J^*/J (FMOD+/+) 8-week-old female mice were obtained from the Jackson Laboratory. The mutant B6(Cg)-*Tyr^c-2J^/J* transgenic FMOD knockout (FMOD-/-) animals were generated previously as described.^14^ The knockout animals were bred, maintained, and housed in pathogen-free conditions at Massachusetts General Hospital, Boston, MA. All animals were kept on 12-hour light-dark cycles under controlled temperatures, and they had free access to standard diet and water.

### Human lupus patient serum

Sera collected from two African American patients with SLE disease activity score (SLE DAI) 8 and 10 and were used in our experiments. Both patients had cutaneous rash and they were on immunosuppressive medications. The two patient sera were equally inoculated among the experimental groups.

Human- and animal-use protocols were approved by the applicable committees of Beth Israel Deaconess Medical Center and Massachusetts General Hospital.

### Injection of human lupus serum

Human lupus serum (100μl) was administered intradermally in the hind leg area of the various mice strains. Twenty-four hours following injection skin was collected for immunohistochemical analysis.

### Scoring of H&E sections

Harvested skin samples were fixed in 4% paraformaldehyde (PF). Healthy skin samples were collected from outside of the injected area. Paraffin embedded sections were cut and stained with Hematoxylin and Eosin (H&E) and histological scoring was done in a blinded manner by a pathologist using morphological criteria published by the International Harmonization of Nomenclature and Diagnostic Criteria (INHAND). The histological sections were analyzed by a grading scheme to evaluate pathologic lesions in the tissues as follows: no lesion (0), minimal (grade 1), mild (grade 2), moderate (grade 3) and marked (grade 4).^17^

### Microscopy and Immunofluorescence staining

H&E-stained sections were imaged by Nikon Eclipse Ti2 microscope. For immunofluorescence staining, the paraffin embedded skin sections were processed as follows: a) rehydration in xylene, ethanol (100%, 95%, 70% and 50%), and phosphate-buffered saline (PBS) solutions; b) blocking; c) incubation with the anti-mouse PE-conjugated Langerin (CD207) (Biolegend) primary antibody overnight; d) staining of nuclei by mounting the slides with VECTASHIELD Antifade mounting medium with DAPI (Vector Laboratories). Fluorescence imaging was carried out with the Nikon Eclipse Ti2 microscope. Captured images were analyzed using the NIS Elements software (Nikon).

## RESULTS

The objective of this study was to investigate the potential contribution of skin pigmentation and melanocyte-secreted FMOD to SLE-associated skin inflammation. It has been established that C57BL/6 mice injected intradermally with serum from patients with SLE develop skin injury^18^. To this end, C57 mice with different fur colors (black vs white) and different expression levels of FMOD (+/+ vs -/-) were injected intradermally with serum of the same lupus patients to induce skin inflammation. Because in previous experiments injection of normal human serum did not induce inflammation,^18^ we did not inject normal serum into our control group. Of Histopathological examination of the skin lesions was performed 24 hours (Fig. 1) after serum injection. Representative sections were hematoxylin & eosin (H&E) stained and evaluated blindly for the following: the presence of epidermal and dermal necrosis with ulceration (0-4), crust formation (0-4), and inflammation in the dermis and subcutis (0-4).

**Figure 1.**
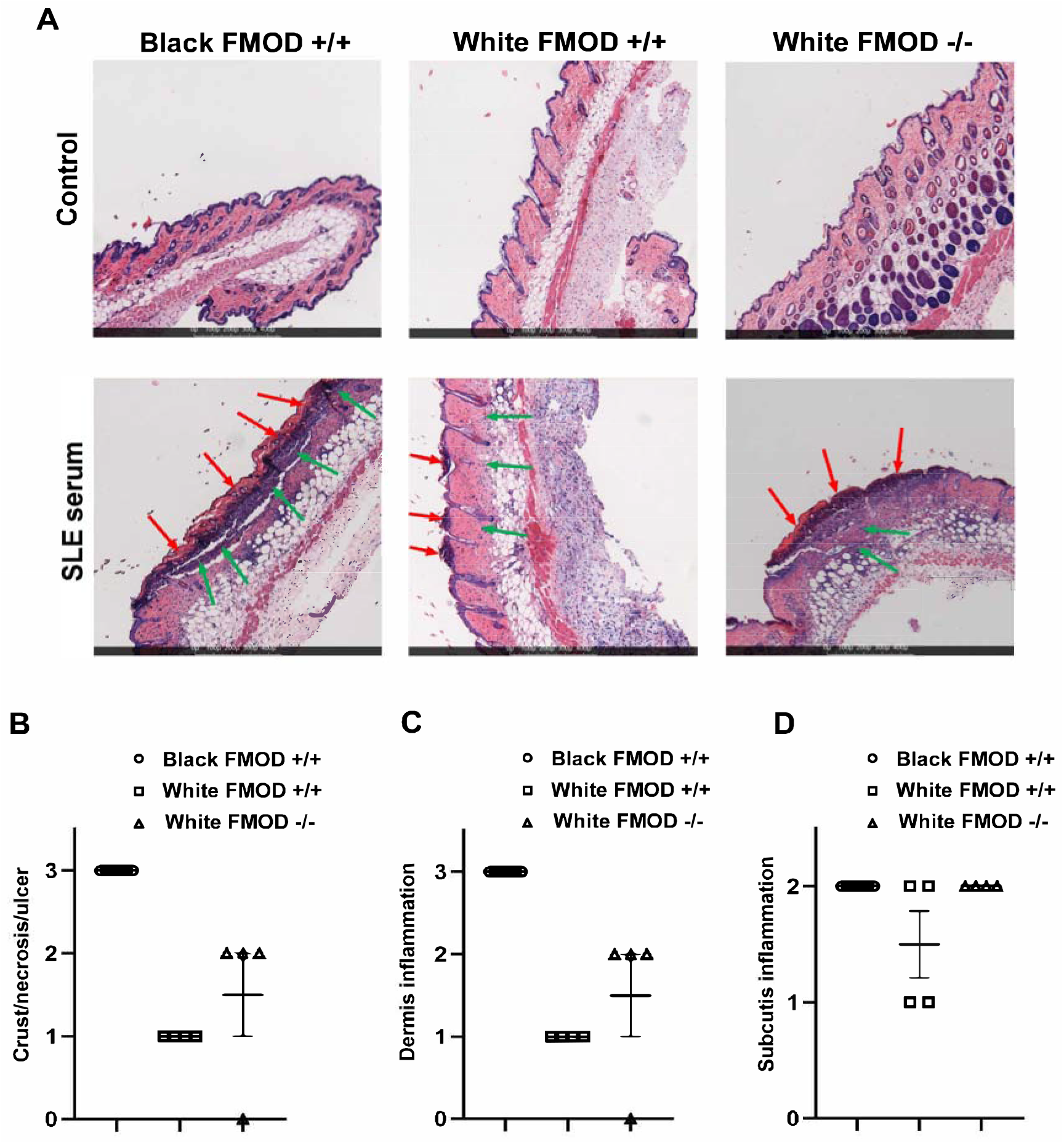
Human lupus serum induces skin inflammation in mice one day after inoculation. **(A)** Black C57 FMOD+/+ (left panel), white C57 FMOD+/+ (middle panel) and white C57 FMOD-/- (right panel) mice were injected intradermally with 100μl serum from lupus patients with disease score 8. Twenty-four hours later mice were sacrificed, and paraffin embedded representative samples were H&E stained. Images were obtained at x10 magnification. **(B-D)** Histological scores for pathological changes including crust/necrosis/ulcer (0-4; red arrows) *(**B**),* dermis inflammation (0-4; green arrows) *(**C**),* and subcutis inflammation (0-4; green arrows) *(**D**)* given in a blinded fashion are plotted.

The histopathological analysis demonstrated that serum from SLE patients induced skin inflammation within 24 hours following injection (Fig. 1A). Quantification of the histological observations revealed that at the 24-hour timepoint the most severe grades of epidermal and dermal necrosis and ulceration, associated with acute dermal inflammation were noted in black C57 FMOD+/+ mice when compared to the white C57 FMOD+/+ mice (Fig. 1B-D), suggesting that dark pigmentation predisposes to the development of more severe inflammation following insult.

Mouse melanocytes isolated from white animals, like human melanocytes isolated from Caucasian people, produce high levels of FMOD when compared to mouse melanocytes isolated from black animals or from black/brown individuals, respectively.^14^ To investigate whether pigmentation associated FMOD plays a role in the skin inflammation observed following serum injection, transgenic FMOD knockout white C57 animals were inoculated with serum from SLE patients. Our findings show that genetic deletion of FMOD resulted in comparable skin injury to black C57 animals characterized by low level of FMOD, suggesting that the presence of FMOD alleviates the gravity of skin lesions associated with SLE serum (Fig. 1).

DCs represent the first line of defense against pathogens entering the body. Increased number of DCs have been noted in inflammatory tissues including the skin in patients with SLE.^19^ Since Langerhans cells are resident epidermal DCs, we assessed the presence of these specialized DCs by immunostaining for Langerin (CD207) on representative healthy (Fig. 2A) and SLE serum-injected skin sections (Fig. 2B). We observed higher expression of Langerin in the skin lesions of black C57 FMOD +/+ and white C57 FMOD -/- mice compared to white C57 FMOD +/+ animals at the 24-time point following serum injection (Fig. 2).

**Figure 2.**
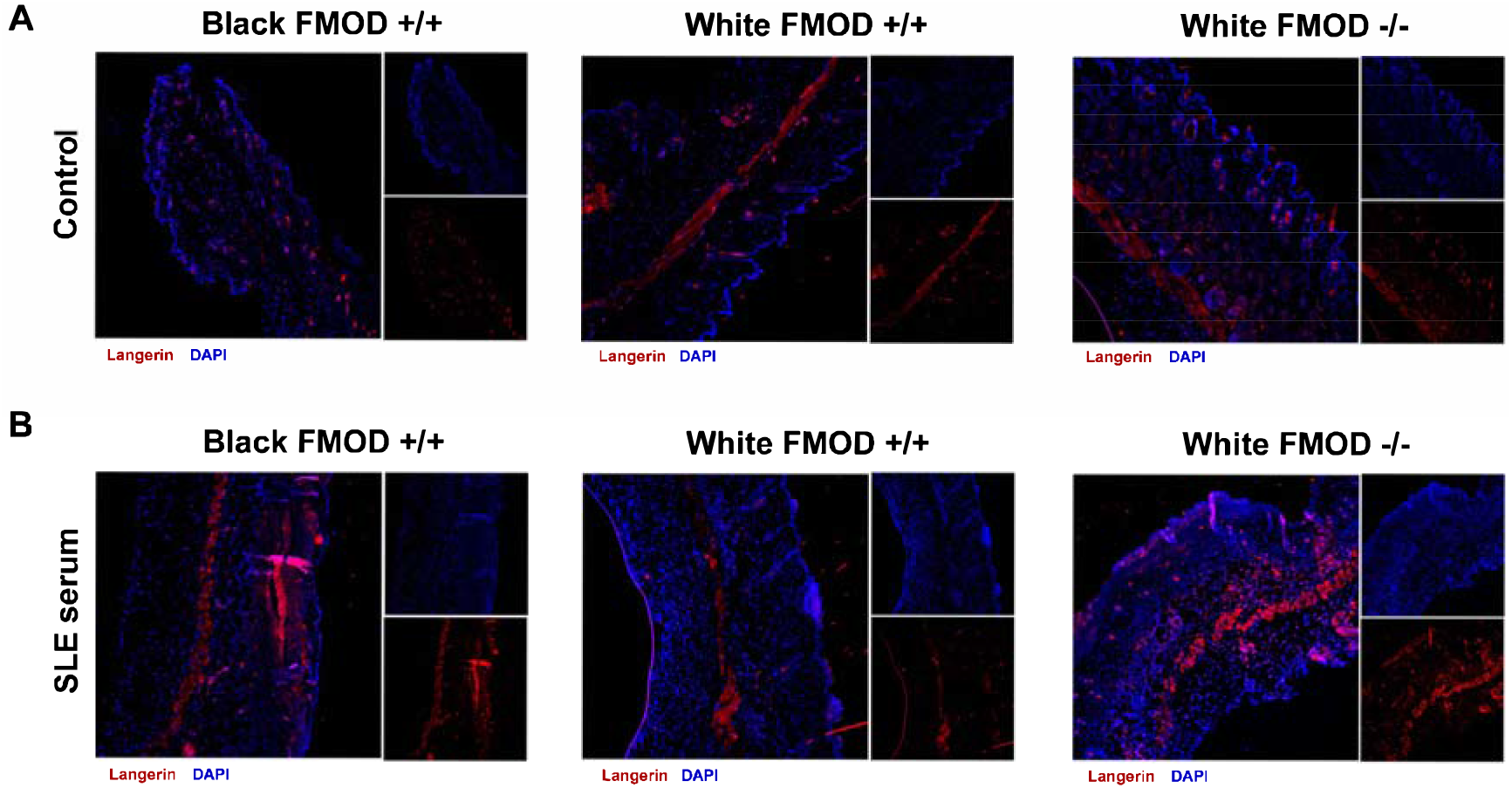
Dendritic cell recruitment is associated with darker pigmentation and low levels of FMOD. **(A, B)** Paraffin embedded sections of healthy and 24-hour human lupus serum injected skin samples collected from Black C57 FMOD+/+ (left panel), white C57 FMOD+/+ (middle panel) and white C57 FMOD-/- (right panel) mice were stained with anti-mouse PE-conjugated Langerin (CD207) antibody. Images were captured at x10 magnifications.

These results suggest that darker skin tones and the absence of FMOD promote the development of more robust inflammation in the skin after injury.

## DISCUSSION

It has been established that race and ethnicity contribute to differences in incidence, prevalence, and severity of inflammation in autoimmune diseases. Specifically, ethnicity is a well-established contributing factor to the severity of both systemic and cutaneous SLE. Understanding the role of racial disparities is critical as it may introduce novel targets and strategies for treatment.

Numerous studies have examined the role of immune cells and cytokines in the initiation and propagation of skin injury. Serum from SLE patients and lupus-prone mice containing IgG are known to induce skin inflammation following local intradermal injection into normal mice. Lupus serum-induced skin inflammation requires the presence of functional TNF-α and TNFR1 and the differentiation of monocytes into dendritic cells (DCs). Activated skin resident DCs enhanced the expression of proinflammatory cytokines and chemokines and enhanced the inflammatory response.^18^ We chose to use the injection of lupus serum to induce skin injury to study the role for melanocytes and FMOD in inflammation in an *in vivo* model system.

We demonstrated that black mice, characterized with highly pigmented melanocytes and low FMOD levels,^14^ develop severe epidermal and dermal necrosis and ulceration, associated with acute dermal inflammation when compared to white mice. More importantly and of mechanistic value, white mice with genetic deletion of FMOD developed a higher histopathological inflammatory response compared to their FMOD+/+ white counterparts. Because we observed higher expression of resident DCs in pigmented FMOD +/+ and white FMOD-/- animals we believe that FMOD protects against skin inflammation by suppressing the expansion of inflammatory DCs. Altogether, this data emphasizes the important link between the production and secretion of FMOD by melanocytes to DC and subsequent T cell activation. Low melanin levels in white strains enable the production of FMOD, which suppresses DC maturation and therefore T cell activation, resulting in reduced tissue inflammation.

This study presents first evidence into the underlying mechanisms of racial disparities in terms of skin inflammation. We provide insight into the involvement of differently pigmented melanocytes and FMOD on inflammation as immunomodulators. Our findings provide cellular and molecular evidence for the development of skin inflammation in dark skinned people and help understand racial disparities along clear mechanistic pathways. Based on our results, we believe that poor FMOD production in individuals with highly pigmented skin tones results in greater activation of DCs, leading to a stronger pro-inflammatory response suggesting the need for changes in treatment strategies to include potential approaches for FMOD production modulation.

## ACKNOWLEDGMENT

This study was supported in part by a grant from NIH (RO1EY024046).

## Notes

### Competing Interest Statement

The authors have declared no competing interest.

